# The N-terminal domain of RfaH plays an active role in protein fold-switching

**DOI:** 10.1101/2021.03.18.436019

**Authors:** Pablo Galaz-Davison, Ernesto A. Román, César A. Ramírez-Sarmiento

## Abstract

The bacterial elongation factor RfaH promotes the expression of virulence factors by specifically binding to RNA polymerases (RNAP) stalled at a DNA signal known as *ops*. This behavior is unlike that of its paralog NusG, the major representative of the protein family to which RfaH belongs. Both proteins have an N-terminal domain (NTD) bearing an RNAP binding site, yet NusG C-terminal domain (CTD) is folded as a β-barrel while RfaH CTD is forming an α-hairpin blocking such site. Upon recognition of the *ops* exposed by RNAP, RfaH is activated via interdomain dissociation and complete CTD structural rearrangement into a β-barrel structurally identical to NusG CTD.

Although RfaH transformation has been extensively characterized computationally, most studies employ tertiary biases towards each native state, hampering the analysis of sequence-encoded interactions on fold-switching. Here, we used Associative Water-mediated Structure and Energy Model (AWSEM) molecular dynamics to characterize the transformation of RfaH, spotlighting the sequence-dependent effects of NTD on CTD fold stabilization. Umbrella sampling simulations guided by native contacts recapitulate the thermodynamic equilibrium experimentally observed for RfaH and its isolated CTD. Temperature refolding simulations of full-length RfaH show a high success towards α-folded CTD, whereas the NTD interferes with βCTD folding, becoming trapped in a β-barrel intermediate. Meanwhile, NusG CTD refolding is unaffected by the presence of RfaH NTD, showing that these NTD-CTD interactions are encoded in RfaH sequence. Altogether, these results suggest that the NTD of RfaH favors the α-folded RfaH by specifically orienting the αCTD upon interdomain binding and also by favoring β-barrel rupture into an intermediate from which fold-switching proceeds.

## Introduction

The NusG/Spt5 family of transcription regulators is universally conserved in all three domains of life. These proteins display a NusG N-terminal domain (NGN) and a number of Kyprides, Ouzounis, Woese (KOW) domains towards their C -terminus [1,2]. The *E. coli* NusG protein displays a single copy of each domain, as in all bacteria and archaea [3]; hence they are normally referred to as N-terminal domain (NTD) and C-terminal domain (CTD) due to their location in the primary structure. The NTD is structurally conserved, folding as an α/β sandwich containing an hydrophobic depression that serves as binding site for the RNA polymerase (RNAP) [4], whereas the CTD folds as a small β-barrel that recruits the ribosome for coupled transcription-translation as well as other partners that regulate transcription [5–7].

The paralogous elongation factor RfaH of *E. coli* is a clear outlier of the NusG family, having an NTD with the canonical NGN structure but a CTD that is folded as an α-helical hairpin rather than the classical KOW β-barrel [8]. This conformation makes up the autoinhibited state of RfaH, as the α-folded CTD is blocking the RNAP binding site located at the NTD, impeding the spontaneous binding to the transcription elongation complex (TEC) that NusG normally undergoes [8]. This autoinhibition is relieved when the transcribing polymerase stalls at a DNA sequence named operon polarity suppressor (*ops*) [9], whose exposed non-template strand forms a hairpin acting as a recruiting partner for RfaH to the RNAP [10–12], promoting interdomain dissociation and NTD binding to the β and β’ subunit of RNAP [13,14]. Strikingly, the dissociated CTD refolds from the initial α-hairpin to a canonical KOW β-barrel which serves as recruiting partner to the ribosomal protein S10, coupling transcription via NTD and translation via CTD [5,13,15].

Numerous studies have addressed the metamorphosis of RfaH through a computational approach, in part due to the difficulties of observing the process in solution since the trigger for RfaH interdomain dissociation is the entire *ops*-stalled TEC. There have been reports indicating the possible pathways through which the isolated CTD may refold from the α-to the β-fold [16–19], which differ from the ones proposed when the CTD is accompanied by the NTD [20–22]. These results suggest that interactions formed between both domains strongly aid in stabilizing the α-fold as well as forming intermediate states that enable the transition between folds [20]. Nevertheless, these studies have focused mostly on the CTD transformation, leaving aside the details of how the NTD stabilizes the α-fold or its effects over the β-folded CTD after the structural transformation occurred. The specifics of NTD-induced energetics on RfaH are not trivial, since the structure of RfaH-NTD [12] displays a more hydrophobic patch than that of NusG [13,23], which has been simultaneously associated to a tighter binding to RNAP, being RfaH NTD the only trigger required for fold-switching back from the active into the autoinhibited state [24].

In this work, we addressed this issue by using the Associative Water-mediated Structure and Energy Model (AWSEM) to determine the effect that the NTD of RfaH has on the overall transformation energetics and the configurational space of both folds. Using umbrella sampling, we determined the change in stability associated to interdomain separation and subsequent fold-switching, recapitulating the experimentally determined equilibrium of the system. Further temperature refolding simulations in the absence of information of known interdomain contacts showed that the highly hydrophobic side of the α-folded CTD consistently looks for an interaction partner and the NTD provides a suitable surface for its stabilization, recapitulating the binding orientation experimentally observed in solved structures of the autoinhibited state of RfaH. At the same time, the NTD interferes with βCTD refolding by mostly trapping it into a β-barrel intermediate, which is also observed in its metamorphic pathway. On the other hand, the β-folded CTD of NusG is impervious to the presence of RfaH NTD, indicating that this folding interference depends on specific interactions encoded in the sequence of both RfaH domains. Altogether, these results suggest that the NTD favors the CTD transformation towards the α-folded CTD by simultaneously stabilizing the α-hairpin and likely disrupting the β-barrel into an intermediate compatible with its refolding pathway.

## Results

### MD simulations of RfaH and its isolated CTD recapitulate their experimental states

The simplest question that can be asked to an energy model about RfaH is whether it can replicate the experimentally observed CTD populations of α and β folds. More precisely, the strong predominance of the α-folded CTD (αCTD hereafter) when RfaH is expressed in its full length in solution [8,12], and the switch of this population to a β-fold (βCTD hereafter) when the CTD is isolated as the result of the NTD-CTD linker being cleaved or by only expressing this domain [5].

To explore this scenario, we set up umbrella sampling simulations that explore the transformation of RfaH for two systems: one in which the full-length RfaH protein is modeled with its CTD in the α-folded (PDB 5ond, αRfaH hereafter) and β-folded state (PDB 6c6s, βRfaH hereafter), and another in which only the CTD of RfaH is modeled in both its αCTD (PDB 5ond) and βCTD (PDB 2lcl) (Fig 1A). Specifically, 50 umbrella simulations were generated for each system, where each one is energetically biased to explore a fraction of the configurations allowed by a reaction coordinate named Q_diff_ (Fig S1). This parameter corresponds to a combination of the individual Q_W_ values of the states to be interpolated, i.e. the fraction of native tertiary contacts [25], resulting in a gradual exploration of decreasing α-specific contacts and increasing β-specific ones (see Methods). This exploration of the transformation was then analyzed using the weighted histogram analysis method (WHAM) [26], and the heat capacity was visually inspected (Fig 1B). From here, to measure the change in stability of α- and β-RfaH, free-energy landscapes were generated at the temperature just before the first peak in heat capacity, as they report the energetics associated to each folded state (Fig 1C and 1D).

**Fig 1.**
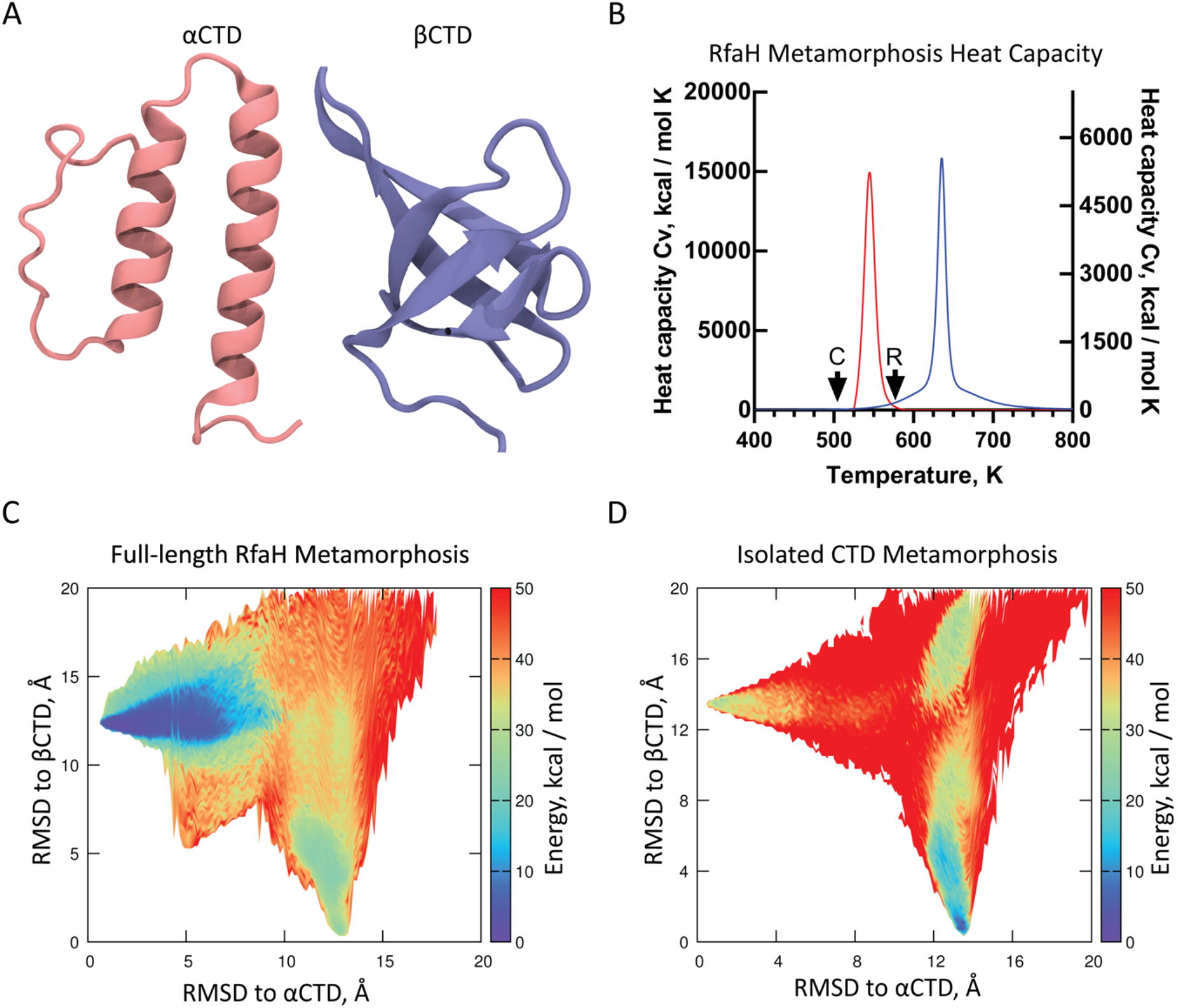
Energetics of the RfaH transformation. (A) Cartoon representation of the two native states of RfaH CTD (residues 110 to 162). (B) Heat capacity profiles of the umbrella sampling simulations of RfaH CTD in the full-length protein (blue) and in isolation CTD (red). The points indicated as C and R mark the temperatures at which the energy landscape was generated for the isolated CTD and full-length RfaH, being 500 and 580 K respectively. (C-D) Energy landscapes for the transformation of RfaH CTD in the full-length protein (C) or the isolated domain (D). The RMSD against the experimental αCTD and βCTD were used as reaction coordinates.

RfaH interdomain contacts were kept as bias for simulating the αRfaH state in the full-length system, while all contacts between the βCTD and NTD in the cryo-EM structure were ignored by increasing the distance in the residue-residue distance matrix beyond the threshold from which a contact is considered to take place (9.5 Å). All the contacts exhibited by the flexible linker in both RfaH initial configurations are also disregarded from the fold-switching simulation as a defined conformation has not been reported for this region.

The results from the umbrella sampling simulations show strikingly different energy landscapes for both RfaH constructs. To observe the preferred folding state, we generated the free energy landscape of RfaH CTD at a temperature just before the first peak in heat capacity (Fig 1B). The full-length protein displays a single, deep energy minimum characterized by a high RMSD against the solved structure of βCTD and a low RMSD towards that of the αCTD (Fig 1C). On the other side of a high energy barrier of ∼50 kcal/mol, βRfaH minimum sits at a higher energy basin than αRfaH (ΔE ∼ 19 kcal/mol at T = 580 K), highlighting the unfavorable nature of the spontaneous transition between both configurations. In contrast, the βCTD is thermodynamically favored over the αCTD (ΔE ∼ 14 kcal/mol at T = 550 K) when this domain is simulated in isolation (Fig 1D). This switch in free-energy difference shows that the αCTD is strongly stabilized only when accompanied by its NTD partner whilst the βCTD is favored in its absence, thus recapitulating the experimental behavior observed for RfaH in solution.

When projecting the two-dimensional free energy profile of the α-to-β transition of RfaH CTD onto a one-dimensional coordinate corresponding to the RMSD difference between the βCTD and αCTD (RMSDβ-α), it can be observed the emergence of a β-intermediate after the destabilization of the native βCTD in the case of the isolated CTD (Fig S2). Also, despite the full-length RfaH is deeply stabilized as αRfaH, the free-energy minima of the βCTD and its intermediate are still observed albeit at a higher energy. Lastly, the native basin for the αCTD in the full-length protein is broad enough to account for fluctuations in the structuredness of the helices, which have been ascertained in both simulations [20,27] and experiments [28]. Alongside the two-dimensional free energy landscapes, our results suggest that the transition route between both RfaH folds using the AWSEM force field is:

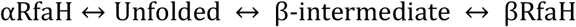

### The NTD of RfaH strongly stabilizes the α-fold and hinders proper βRfaH refolding

One disadvantage of the umbrella sampling simulations is that, by directly employing the number of native contacts of the system in αRfaH and βRfaH as collective variables to drive the structural interconversion of RfaH, the formation or disruption of interdomain contacts between specific residue pairs is also biased. As a consequence, one is unable to directly evaluate how the appropriate binding configuration between the NTD and CTD is encoded within RfaH sequence. However, AWSEM allows to restrict the use of structural biases only towards local-in-sequence interactions by providing a “fragment memory” potential (see Methods) that limits the configurational exploration of short segments of the protein to those of a reference structure [29]. By not providing information about contacts between the NTD and CTD, these simulations freely explore the interdomain interaction landscape. A similar simulation strategy has been previously employed to correctly predict the binding interfaces of both homodimers and heterodimers [30].

Using a temperature gradient through long molecular dynamics simulations (3·10^7^ timesteps of 5 fs, compared to previously reported folding annealing simulations of 4·10^6^ timesteps [29] and 6·10^6^ timesteps[31]), 100 models with a single fragment-based memory were allowed to refold starting from random unfolded conformations (Q_W_ < 0.1) (Table S1). In these single-memory models only the NTD and CTD of RfaH, but not the linker connecting both domains, were given fragment memories, and these memories are withdrawn from a single reference structure, either αRfaH or βRfaH. This approach leaves the linker that connects both domains with a major conformational freedom (Fig 2B) and results in the C- and N-terminal domains being structurally uncoupled, as the 10-residue long connector that exist between them is not part of the structural bias and therefore disrupts memory continuity. Therefore, these folding simulations emulate the solution behavior expected for a two-domain protein.

**Fig 2.**
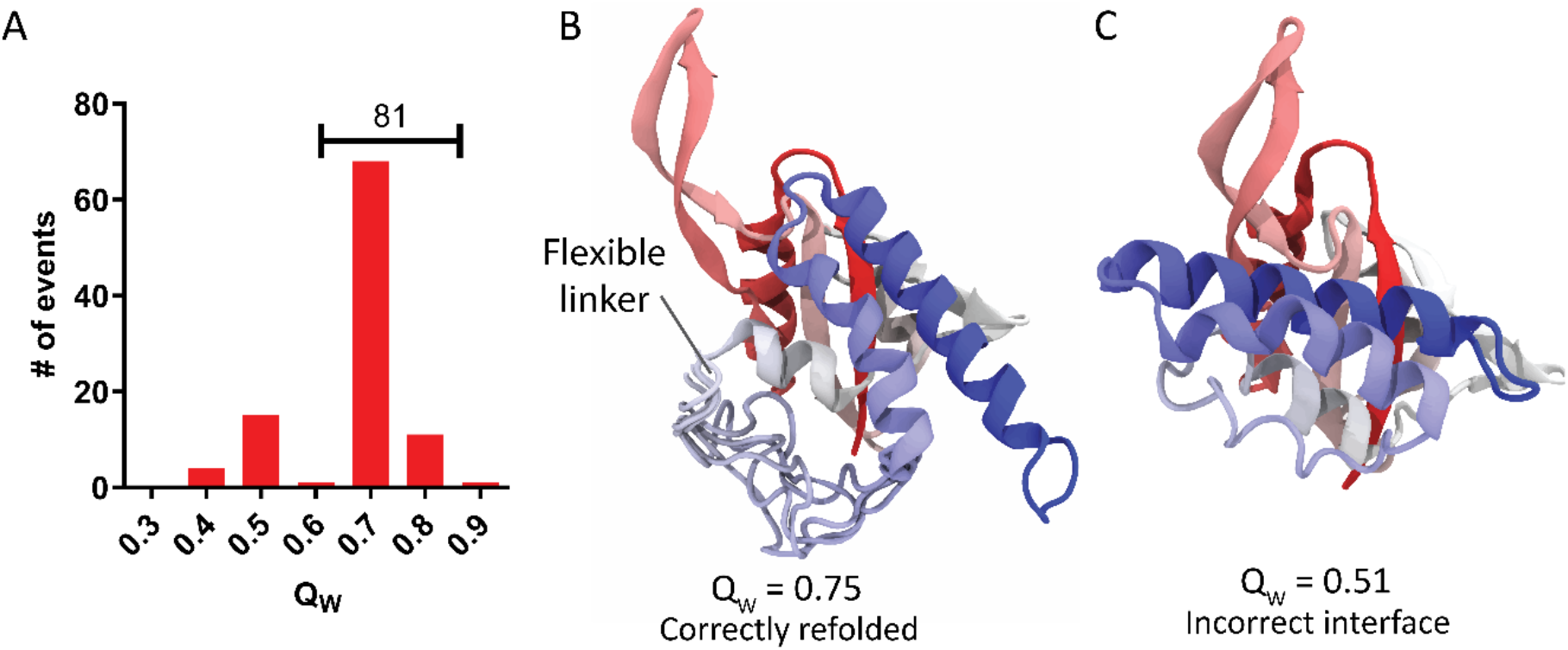
Refolding efficiency of αRfaH. (A) Distribution of tertiary contacts (Q_W_) in the final structure of the 100 refolding simulations generated for αRfaH using a single memory (PDB 5ond). (B-C) Representative final structures after αRfaH refolding with high (B) and low (C) Q_W_ respectively.

Refolding simulations employing αRfaH as the single memory reference structure (PDB 5ond) show that nearly 81% of the trajectories arrive at the native state (Q_W_ = 0.75, Fig 2A). These predicted structures are characterized by the proper orientation and binding of the αCTD against the interdomain region of the NTD (Fig 2B), recapitulating the experimentally solved structure of RfaH in its autoinhibited state [8]. This specificity is achieved despite the lack of structural biases on the interdomain interface and linker regions, and thus a result of sequence determinants in both the NTD and CTD of RfaH encoding this behavior. In fact, the linker is not stabilized in a particular conformation (Fig 2B) and does not form stable contacts with any domain. In all other trajectories the interdomain interface is formed incorrectly, although both the NTD and αCTD reach their native conformations mostly due to the fragment memory bias (Fig 2C). These finding suggest that the NTD-side of the interdomain interface is strongly guiding both the binding orientation and contact formation between NTD and αCTD.

To further assess the effect of the NTD hydrophobic patch in CTD folding, the same refolding experiment was performed for βRfaH extracted from the cryo-EM structure (PDB 6c6s). To enlighten the effect that the NTD could have on βCTD refolding, the resulting structures are compared with equivalent refolding simulations on the isolated βCTD (PDB 2lcl). The results of βRfaH and βCTD refolding experiments are summarized in Fig 3. For the isolated CTD, the βCTD refolds with an efficiency similar to that of αRfaH (75%), with the remainder of the simulations reaching an intermediate state characterized by a lower Q_W_, in which only the three larger β-strands of the barrel are folded (Fig 3B). In contrast, the presence of RfaH NTD reduces the CTD refolding efficiency to only 29%, whereas all other refolding trajectories become trapped in the same β-intermediate observed for the isolated βCTD. These results suggest that the stabilization of this intermediate is a result of specific NTD-CTD interactions stablished during the folding process of βRfaH.

**Fig 3.**
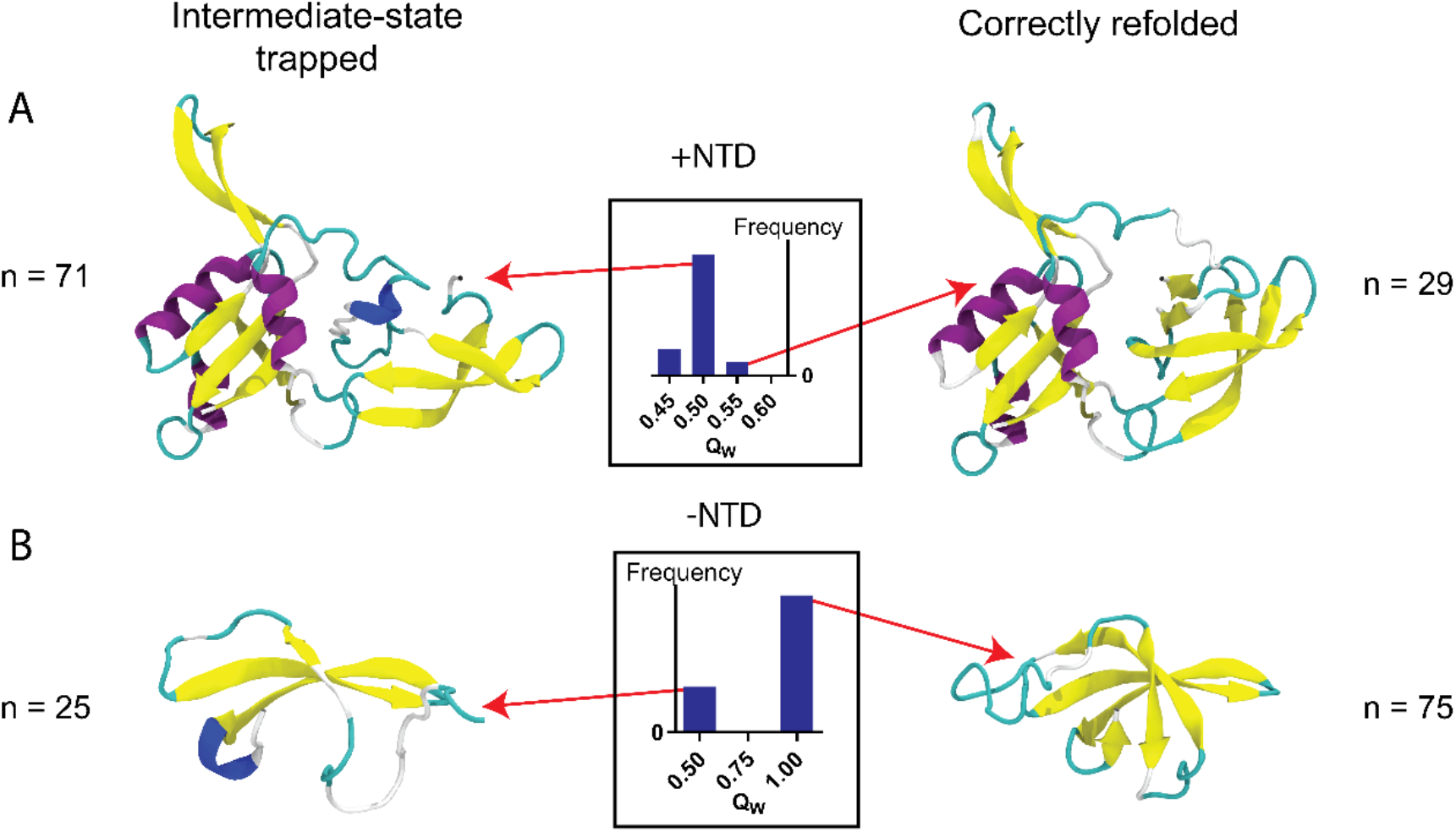
Refolding of βCTD in context of the full-length βRfaH and in isolation. Representative final structures after βCTD refolding in the context of the full-length protein (single memory, PDB 6c6s) (A) and in isolation (single memory, PDB 2lcl) (B). The histograms represent the Q_W_ distribution of the final structures. The intermediate state is formed by the three largest β-strands that form the CTD barrel, namely strands β2, β3 and β4. For further details of the refolding statistics, please refer to Table S1.

To determine that the βCTD intermediate is stabilized by specific interactions between both RfaH domains, a harmonic potential was used to maintain the NTD and CTD domains away from each other during refolding simulations of βRfaH. Upon keeping both domains apart throughout the simulation, the βCTD mostly refolds as if it was isolated, with 66% of cases achieving complete refolding (Table S1). Also, two additional systems were used for refolding simulations: i) the isolated CTD of NusG (PDB 2jvv), a protein that shares almost identical secondary and tertiary structure but lacks any observable metamorphic feature, and ii) a chimeric protein connecting the NTD of RfaH with the CTD of NusG, in which it is expected that no specific interdomain interactions are formed given the divergent evolution of RfaH and NusG [32]. Remarkably, when the isolated CTD of NusG (PDB 2jvv) and its fusion to RfaH NTD (PDB 5ond) were used as input for refolding simulations, the totality of the simulations reached the β-folded state of NusG CTD, regardless of the presence of the NTD (Fig S3, Table S1). Although NusG CTD also traverses through a three-strand intermediate state during refolding, it does so at a slightly higher temperature than the βCTD of RfaH (Fig S4). Altogether, these data strongly suggest that an interruption in the β-barrel folding process is caused by specific interactions stablished between RfaH domains.

To gain insights about the interdomain interactions that stabilize the folding intermediate of RfaH βCTD, we determined the per-residue tertiary contacts that are minimally or highly frustrated using the protein frustratometer [33]. For this end, a representative structure of the most populated cluster of the intermediate-trapped or completely refolded βCTD, both in isolation and in the context of full-length RfaH, were analyzed using the web version of the protein frustratometer (http://frustratometer.qb.fcen.uba.ar). For the completely refolded βCTD, a similar amount of highly and minimally frustrated contacts is observed for each CTD residue, regardless of the presence of the NTD (Fig 4), with the exception of residues 119-126 and 145, which greatly increase the amount of minimally frustrated contacts when refolded in the full-length RfaH. These sets of residues have been identified to be relevant for the stability of the βCTD in previous simulations using dual-basin structure-based models [20] and also for the stability of the autoinhibited state of RfaH in recent NMR experiments of the transformation of RfaH [34]. In stark contrast, the β-barrel intermediate of the CTD forms more minimally frustrated contacts when in the presence of the NTD than in isolation (Fig 4), particularly doubling the number of these type of contacts in the region corresponding to strand β_1_ and the loop preceding strand β_2_ (residues 114-123). Despite not forming the strand β_1_, such region becomes highly stabilized by bridging interactions between the NTD and the β-barrel intermediate and serves as the interface between the two domains.

**Fig 4.**
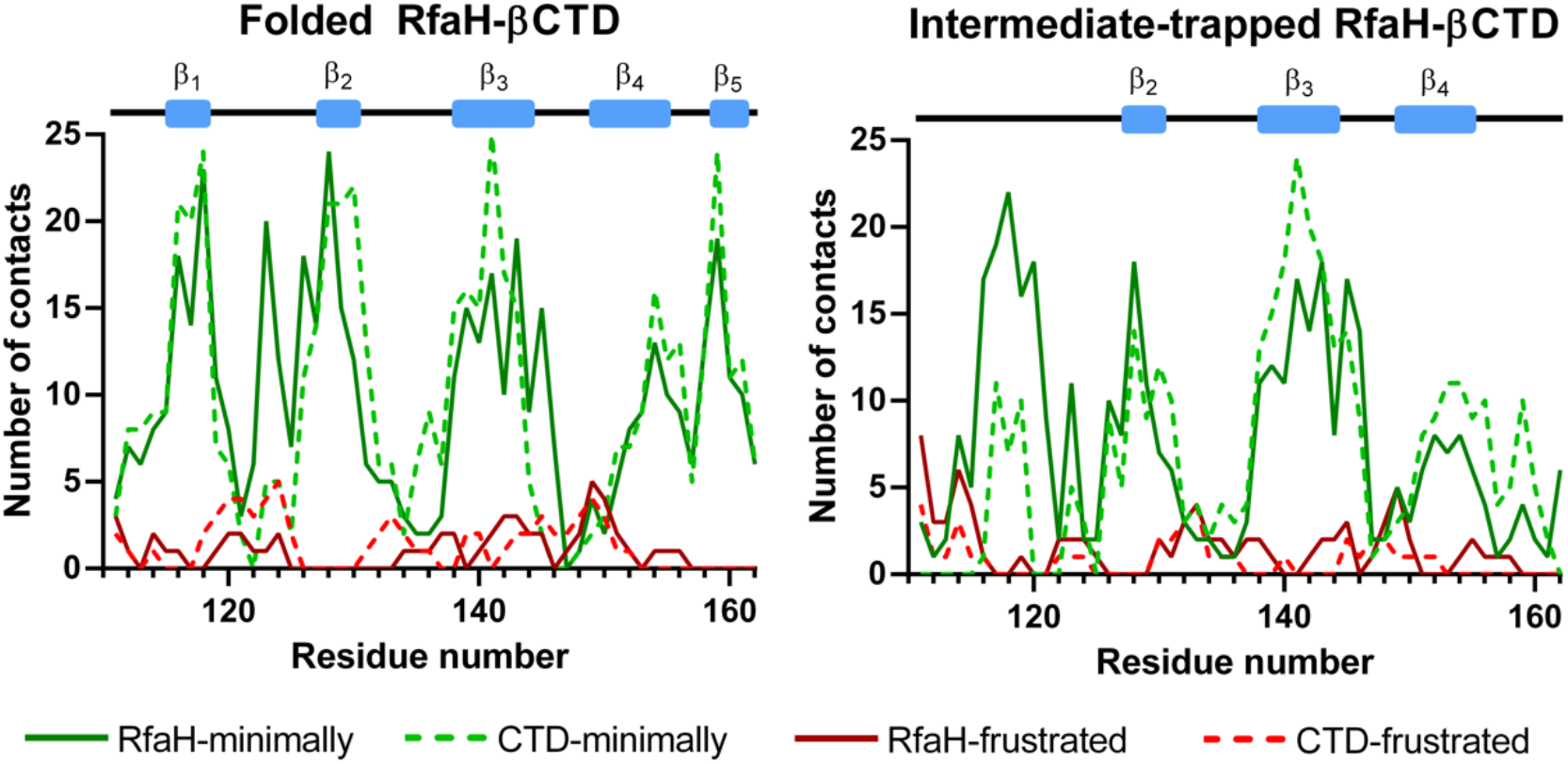
Frustration analysis of βCTD and its folding intermediate in both full-length RfaH and in isolation. The number of highly (red) and minimally frustrated (green) contacts is shown for the CTD in isolation (dashed line) and in the context of full-length RfaH (solid line).

## Discussion

*E. coli* RfaH is known as one of the most dramatic examples of protein fold-switching. In solution, RfaH folds into an autoinhibited state in which the αCTD tightly binds to the NTD, whereas its active state, in which the NTD and βCTD fold and appear to behave independently, is thermodynamically unfeasible in the absence of *ops*-stalled TEC [13]. In contrast, the non-metamorphic *E. coli* NusG only transiently forms interdomain interactions and exists in solution as a two-domain protein [5,23]. Our simulations using the AWSEM molecular dynamics and force field package correctly model RfaH in all its conformations and recapitulate its behavior in solution, evidenced as the switching of the energetic minimum between αCTD and βCTD when breaking interdomain interactions that has also been observed in previous computational works on full-length RfaH using various simulation strategies [20,27].

More importantly, our refolding simulations show that the number of trajectories that successfully reach the β-folded CTD in the context of full-length RfaH is a minority when compared to the cases in which the CTD becomes trapped in a three-strand β-barrel intermediate, and almost three times less successful than refolding of αRfaH. We also demonstrate that a significant number of minimally frustrated NTD-CTD interactions, some of which involve residues that also participate in interdomain interactions taking place in the autoinhibited state of RfaH, interfere with proper β-fold formation by stabilizing its intermediate state. These results suggest that the thermodynamic stability of the autoinhibited state of RfaH is not only due to the compatibility between the αCTD and NTD but also due to an active destabilization of the β-barrel by the NTD, which increases the probability of the β-state being trapped in this folding intermediate. Moreover, while refolding of the CTD of the RfaH paralog NusG successfully reaches the β-folded state, the transient observation of a structurally similar intermediate state also suggests that it is the nature of the NTD and CTD sequence of RfaH that drives the interdomain interaction and ultimate trapping into this state.

Multiple reports have studied the metamorphic process of RfaH CTD in the context of the isolated domain [16–19] and the full-length protein [20–22], but only a few have described the β-intermediate observed here during βCTD refolding. One of such works corresponds to the computational study of the α-to-β transition of the isolated CTD of RfaH through targeted molecular dynamics and Markov state models using an adaptive seeding method, in which several en-route ensembles collectively suggests that strands β2, β3 and β4 are relatively stable and form earlier during refolding towards the β-state [16]. Additionally, our previous work with full-length RfaH using dual-basin structure-based models also identified a βCTD-like intermediate that is either free or interacting with the NTD, but with a different topology [20]. Lastly, recent unbiased explicit solvent simulations of the spontaneous α-to-β fold-switch of RfaH CTD using a replica exchange with hybrid tempering method exhibits three-stranded and four-stranded intermediates before reaching the β-folded CTD [35]. Nevertheless, none of these works described the active role of the NTD in stabilizing such intermediate state nor characterized its role as part of the β-barrel folding process.

We believe that this intermediate and its NTD-dependent stabilization has been overlooked due to either the granularity of the model used, the absence of sequence-dependent potentials or the velocity with which the system is being driven out of the equilibrium. In fact, the sequence-dependent potential embedded on AWSEM shows its capabilities when simulating the correct refolding of αRfaH to a high fraction of native contacts Q_W_ even in the absence of knowledge-based contact information of the interdomain interface and the linker connecting both domains, meaning that these simulations are robust enough to discriminate the interactions arising from RfaH sequence in terms of NTD-CTD association. The observation that NusG CTD, unlike RfaH βCTD, is not affected by RfaH NTD in these simulations is confirmation of the latter.

These arguments, alongside the observation of this intermediate in both NusG and RfaH βCTD folding pathways, also suggest that this intermediate is a topological solution to the small β-barrel folding process, which could also be necessary for the transition between the α- and β-folds of RfaH. While our previous work using hydrogen-deuterium exchange mass spectrometry show no apparent differences between NusG CTD and RfaH CTD and no indications of intermediate states under native conditions [28], it is possible that the intermediate state observed here requires the addition of chaotropic agents to favor its abundance. It can be presumed that the destabilization of the native state using such approaches not only would favor the intermediate population but also the unfolded state.

All in all, our simulations indicate that the NTD actively participates in thermodynamically favoring the autoinhibited α-state by properly orienting the αCTD and correctly specifying the interactions occurring upon interdomain interface formation and also by destabilizing the β-folded CTD into a folding intermediate. Such intermediate could be potentially observed by studying the equilibrium unfolding of the isolated CTD, as it was observed here during the refolding process of the isolated CTD of RfaH and NusG as well as part of the metamorphic pathway in full-length RfaH. We also hypothesize that this βCTD destabilization by the NTD is the initial step for RfaH to fold-switch back into the autoinhibited state. This idea is compatible with the observation of RfaH stably binding the ribosomal protein S10 through its βCTD when bound to the TEC [34], as in such state the NTD hydrophobic patch is blocked by RNAP. Therefore, the effect of the NTD over the βCTD can only be observed when releasing the active state of RfaH from the TEC.

## Methods

### The AWSEM molecular dynamics model

The Associative Water-mediated Structure and Energy Model, AWSEM, [29] is a C_β_ granularity model implemented in LAMMPS [36]. This model contains five energy terms, which are [29]:

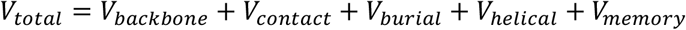

Of these terms, the backbone energy term guides the atoms to a protein-like geometry, while the contact term is responsible for the formation of residue-residue interactions in an amino acid-dependent manner. The burial energy term regulates the exposure to the solvent of the protein core, depending on the residue identity. The helical term replicates the hydrogen bonding pattern of carbonyl oxygen-amide nitrogen formed in α-helices depending on the involved residues helical propensity. Finally, the memory term is a local bias applied to overlapping fragments from 3 to 9 residues that guides C_α_ and C_β_ distances to those of a reference structure. This potential can be guided to multiple structures, or as used in this work, limited to a single reference structure [29]. The λ_FM_ used in this work is of 0.3 compared to the default 0.2, resulting in a higher cooperativity observed in protein refolding experiments.

### Calculation of Q_diff_ and umbrella sampling

For the umbrella sampling method, 50 simulations of 2.4·10^7^ timesteps were run and energy and trajectory frames were collected every 1,000 timesteps. The initial configuration was that of the unfolded isolated CTD (dual memories: 2lcl for βCTD and 5ond for αCTD) or unfolded CTD in the context of full-length RfaH (dual memories: 6c6s for βRfaH and 5ond for αRfaH). The simulations sampled fractions of an order parameter called Q_diff_ which corresponds to [25,37]:

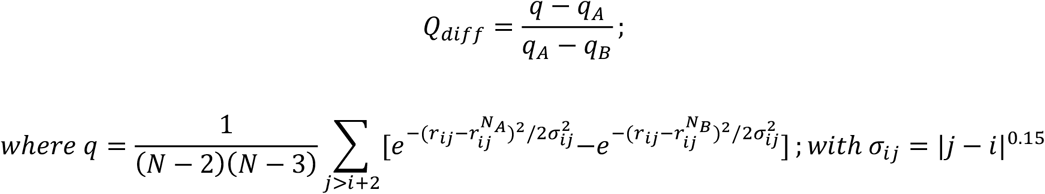

Where *q*_A_ and *q*_B_ are constants obtained by evaluating the *q* function in the two structures to which the transition is to be interpolated, and *r*_ij_ measures the C_α_ distance between residue *i* and *j*. This is evaluated for all contacts between j>i+2 residues whose C_α_ are at a distance of

9.5 Å or below. Using the Q_diff_ value, a bias is applied by adding a new potential to the system with the form:

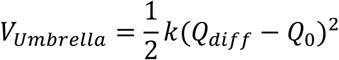

Where *k* is the harmonic potential constant, here 1,500 kcal·mol^-1^, and Q_0_ being the center of the distribution of a Q_diff_ value ranging from 0.02 to 1.00 by increments of 0.02. From these simulations the potential energy and Qdiff values were obtained for each frame as well as the C_α_ RMSD of the best fit against both CTD native folds, which was calculated using VMD [38]. The RMSD against αCTD and βCTD were then used as reaction coordinates for thermodynamic analysis using the WHAM algorithm [26] implemented in Java [39]. For this analysis, the first 3,000 frames were excluded as this was the equilibration frame from the unfolded state to the desired biased configuration.

### Refolding simulations

For these simulations, random initial unfolded configurations for each system were generated by running 100,000 timesteps of a simulation without any potential but the backbone energy term, saving a simulation restart configuration every 10,000 timesteps. The restart configuration with the lowest Q_W_ value, which in all cases was below 0.1, was used as a starting configuration for the refolding simulations. All 100 simulations were randomly assigned initial velocities and run for 3·10^7^ timesteps of 5 fs, during which the temperature linearly decreased from 1000 K to 400 K. All constructs were completely unfolded at the initial temperature (as in Fig S4) and either completely refolded, trapped into an intermediate state or misfolded at the final temperature (Table S1). The final structures of these simulations were clustered by calculating pairwise best-fit RMSD [40] using Chimera [41]. For the representative member of each cluster, as well as for non-clustered models, the secondary structure assignment was calculated using STRIDE [42]. These secondary structure assignments are summarized in Table S1 alongside the corresponding Q_W_, which is a measure of structural similarity to a given structure and obtained using the formula:

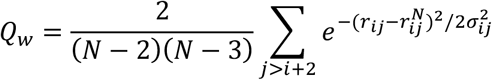

Where, similarly as for Q_diff_, *r*_ij_ measures the C_α_ distance between residues *i* and *j* for the current or reference (superscript N) structure. Via visual inspection of the members of each cluster and individual models, misfolding of the NTD was identified where possible and also reported.

## Supporting Information

Table S1 and Fig S1-S4 are available in the Supporting Information file.

## Data availability

All produced trajectories and data analysis, as well as the AWSEM and LAMMPS version used for these simulations, will be available for download at the laboratory’s website (https://pb3.sitios.ing.uc.cl). The remainder of the data is shared upon reasonable request.

## Author Contributions

P.G.-D., E.A.R. and C.A.R.-S. designed the research, P.G.-D. and E.A.R. performed the simulations, P.G.-D., E.A.R. and C.A.R.-S. analyzed the data, P.G.-D., E.A.R. and C.A.R.-S. wrote the manuscript, E.A.R. and C.A.R.-S. were leading investigators.

## Acknowledgements

This research was funded by the National Agency for Research and Development (ANID) FONDECYT Regular Grant 1201684. PGD was supported by ANID Doctoral Scholarship 21181705 and a Red MacroUniversidades award for graduate mobility to Universidad de Buenos Aires. EAR has a research grant from Agencia Nacional de Promoción Científica y Tecnológica (PICT 2016-0014).

